# Functional heterogeneity of lymphocytic patterns in primary melanoma dissected through single-cell multiplexing

**DOI:** 10.1101/409011

**Authors:** Francesca Maria Bosisio, Asier Antoranz, Yannick van Herck, Maddalena Maria Bolognesi, Lukas Marcelis, Clizia Chinello, Jasper Wouters, Fulvio Magni, Leonidas Alexopoulos, Marguerite Stas, Veerle Boecxstaens, Oliver Bechter, Giorgio Cattoretti, Joost van den Oord

## Abstract

In melanoma, the lymphocytic infiltrate is a prognostic parameter classified morphologically into “brisk”, “non-brisk” and “absent” entailing a functional association that has never been proved. Recently, it has been shown that lymphocytic populations can be very heterogeneous, and that anti-PD-1 immunotherapy supports activated T cells. Here, we characterize the immune landscape in primary melanoma by high-dimensional single cell multiplex analysis in tissue sections (MILAN technique) followed by image analysis, RT-PCR and shotgun proteomics. We observed that the brisk and non-brisk patterns are heterogeneous functional categories that can be further sub-classified into active, transitional or exhausted. The classification of primary melanomas based on the functional paradigm also shows correlation with spontaneous regression, and an improved prognostic value than that of the brisk classification. Finally, the main inflammatory cell subpopulations that are present in the microenvironment associated with activation and exhaustion and their spatial relationships are described using neighbourhood analysis.

## Introduction

The lymphocytic infiltrate in melanoma is a prognostic parameter reported by the pathologist as patterns of tumour-infiltrating lymphocytes (TILs). The “brisk” pattern (diffuse or complete peripheral TILs infiltration) has a better prognosis compared to the “non-brisk” (tumour areas with TILs alternate with areas without TILs) or to the “absent” pattern (no TILs or no contact with melanoma cells)^1,2,3^. There are multiple pitfalls in the purely morphological evaluation of TILs^4^, but the most important one is that morphology alone cannot determine their activation status. The meaning of the word “brisk” according to the dictionary is “active, energetic”, a definition implying a functional connotation starting from a morphological evaluation. Surprisingly, this functional connotation has never been proved. The contact between cytotoxic lymphocytes (Tcy) and melanoma does not always lead to tumour eradication but, due to immune modulation, can also result in Tcy inactivation. These “exhausted” Tcy would still be morphologically present, indistinguishable from active lymphocytes. Moreover, the morphological^5^ and functional^6^ side of the tumour microenvironment has been separately investigated, but integration of both types of data is still lacking.

In recent years, several methods for single cell-analysis have been implemented in order to obtain a high-resolution landscape of the tumour microenvironment^7^. According to a recent review^8^, high resolution means to characterize not only the immune infiltrate but also to define the spatial distribution of each component within it, which allows one to make inferences about cell-cell interactions. Nevertheless, most of these methods rely on dissociation of the cells from fresh material, an impractical option in primary melanomas, nowadays diagnosed at an early stage, with very limited material, and resulting in loss of the knowledge of the spatial distribution of each component within it. Moreover, several studies for predictive biomarkers relies on peripheral blood^9^. Though, the functional status of circulating inflammatory cells can completely change entering the tumour site^10^. Intuitively, it is the behaviour of the inflammatory cells in the surroundings of the tumour that will make a difference in terms of prognosis and response to therapy.

Here, we characterize the immune landscape at single cell-level in primary melanoma based on a panel of 39 immune markers applied on one single tissue section through a high-dimensional multiplexing method^11^, an RT-PCR expression evaluation and a shotgun proteomic analysis. This approach allowed us a) to further categorize the brisk and non-brisk morphological patterns of TILs into three functional categories; b) to define the correlation between T-cell activation and spontaneous melanoma regression; c) to investigate the most important inflammatory subpopulations involved in TILs exhaustion (Figure 1).

**Figure 1.**
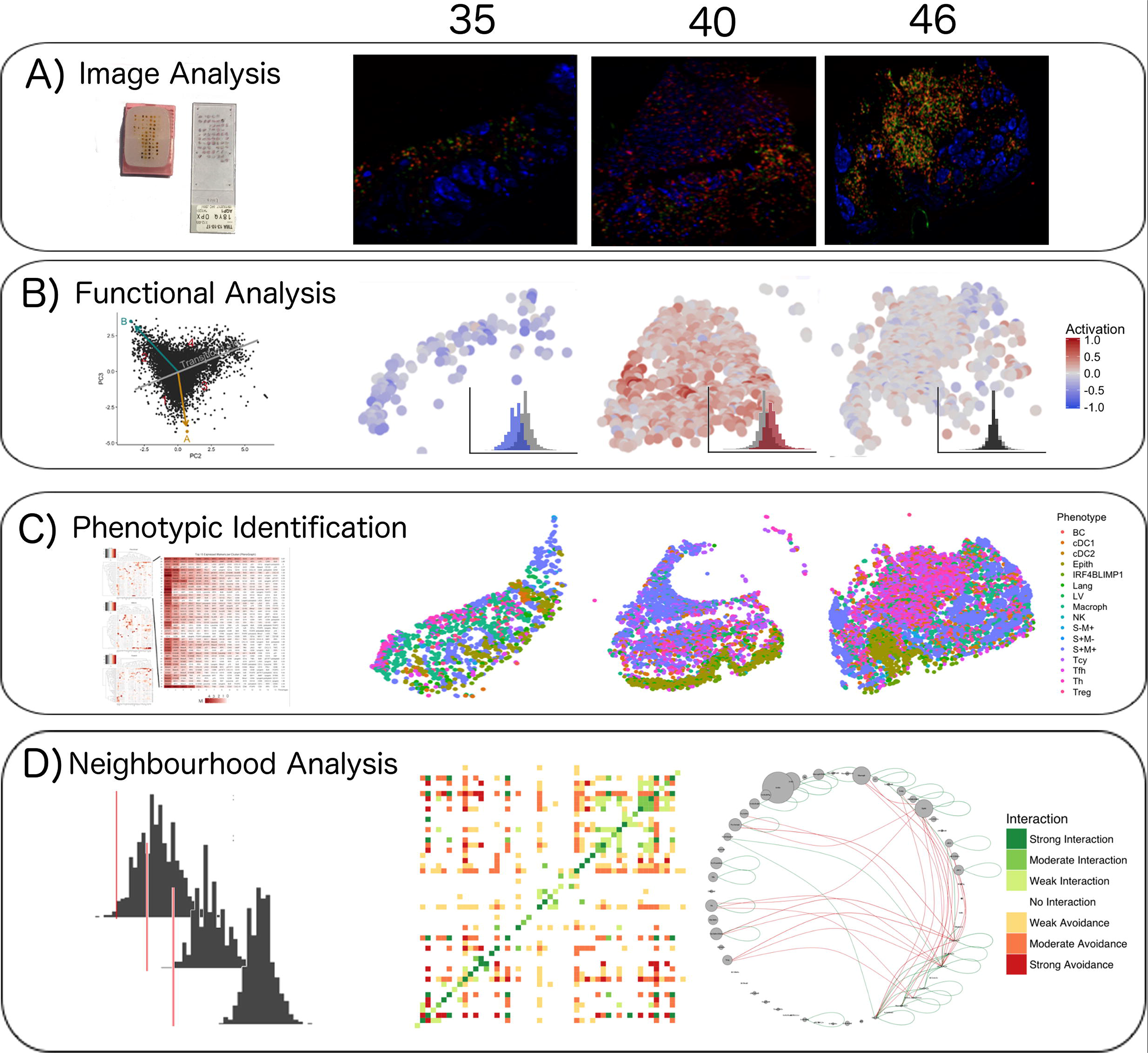
Single cell analysis scheme. A) 60 cores were obtained from 29 patients, assembled in a Tissue Micro Array, and analysed using the MILAN technique^11^; B) CD8+ cells were analysed using image analysis for functionality using an activation parameter derived from multiple activation and exhaustion markers evaluated at single cell resolution; C) All the cell populations in the cores were phenotypically identified using consensus between three clustering methods and manual annotation from the pathologists; D) The social network of the cells was analysed using a permutation test for neighbourhood analysis in order to make inferences on cell-cell interactions. All steps are exemplified using three representative images (35, 40, and 46).

## Results

### Functional analysis of TILs

We classified each CD8+ cell in the TMA cores as part of a spectrum (“functional status”) ranging from “active” (CD69high and/or OX40high) to “exhausted” (TIM3highCD69lowOX40low) (Figure 2 and Supplementary Data 1). The assignation of a functional status to each core in the TMA (“core status”, see methods) yielded 17/60 cores defined as active, 23/60 in transition, and 20/60 defined as exhausted. Core classification allowed the assessment of the heterogeneity of the immune response in different areas of the melanoma for the same patient. From the 29 patients included in the analysis, 8 patients allowed to sample only a single core due to the size of the melanoma and could not be included in this analysis. From the 21 remaining patients, 10/21 showed homogeneous core statuses: 4 active, 5 in transition, and 1 exhausted; and 11/21 showed heterogeneity. Correlation with clinical survival (overall survival, OS) showed that patient classification based on functional status has an improved prognostic performance (log-rank p.value = 0.079) when compared with the brisk morphological classification (log-rank p.value = 0.36) (Figure 2.E).

**Figure 2.**
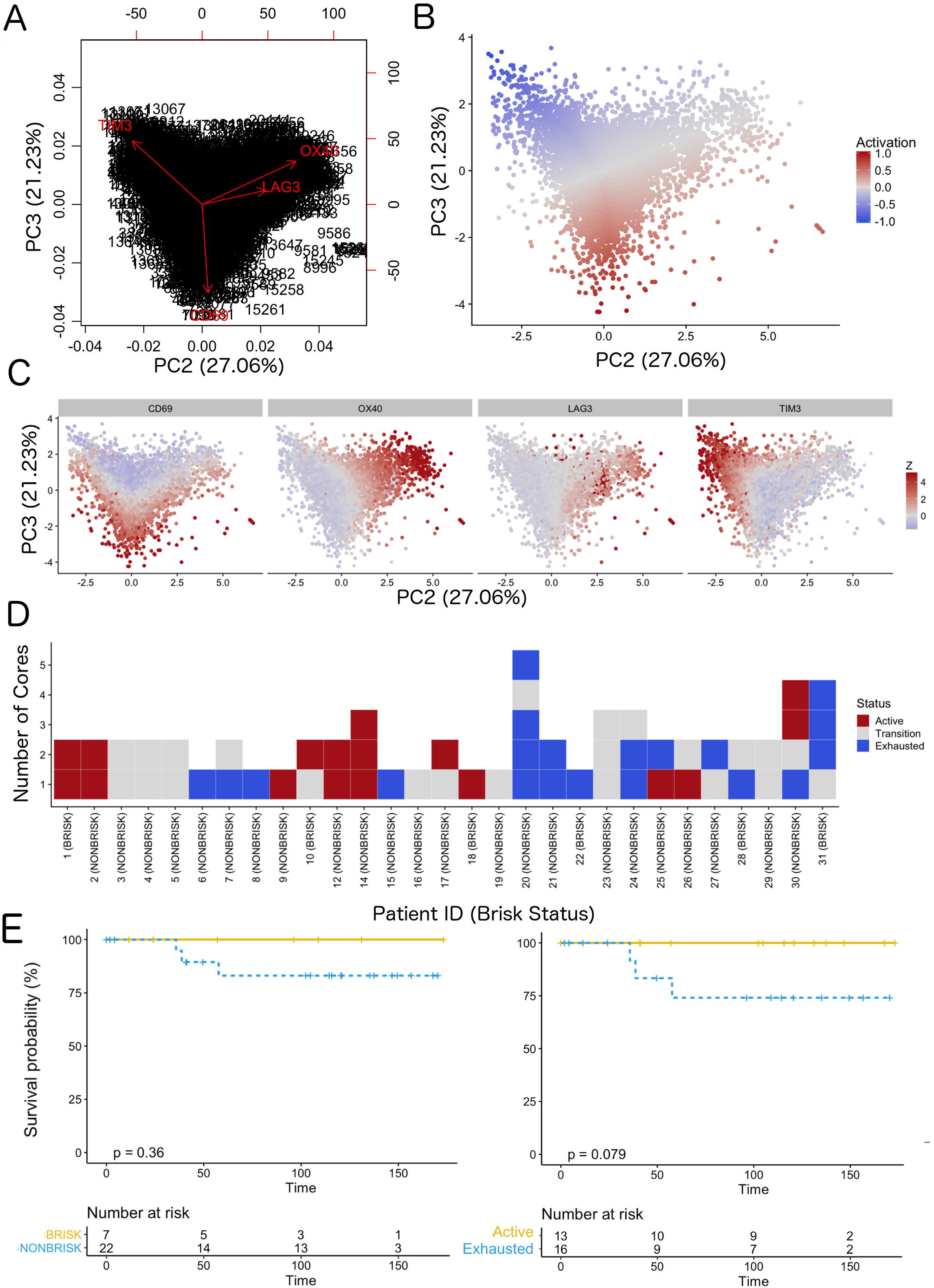
Definition of activation and implications in overall survival. A biplot showing the projection of the cells as well as the rotation vectors of the markers over PC2 and PC3 (A) was the first step to define a gradient of activation going from the maximum projected value of CD69 (maximum activation) to the maximum projected value of TIM3 (maximum exhaustion) (B). C) Z-scores of the original markers over PC2 and PC3. D) Visual representation of the inter- and intra-patient heterogeneity. Each core is assigned an activation status (‘Active’, ‘Transition’, or ‘Exhausted’). The cores are grouped for each patient, giving an at-a-glance representation of the heterogeneity of the activation status in different areas of the melanoma in the same patient. E) The functional definition of activation/exhaustion shows improved prognostic value when compared to the brisk morphological parameter (p.value = 0.075 vs p.value = 0.31 log-rank test).

We then checked whether the core status was significantly associated with spontaneous regression of the tumour, regarded as the result of a successful Tcy immune response, and with other histopathological parameters (histological subtype, ulceration, Breslow thickness, mitoses). Late regression areas indeed showed significant differences in the mean level of activation of the cores as compared to early regression (p = 0.022) and no regression (p = 0.031). No significant differences were instead found between early regression and no regression. Higher levels of activation were found in Lentigo Maligna Melanoma (p = 0.02). However, since only three LMM cores from two patients were included in our data set, no definite conclusions can be drawn from this data. The other histopathological prognostic parameters did not show significance (Table 1).

**Table 1.** Histopathological correlation. Histopathological parameters were correlated with core status (Active/Transition/Exhausted) using pairwise t-tests with pooled standard deviation.

### Phenotypic identification

The inflammatory subpopulations were identified using three different unsupervised clustering methods (KMeans, PhenoGraph, and ClusterX^12^) followed by manual annotation of the clusters by an expert pathologist (FMB). Cells were evaluated for consistent cell phenotype as described in the methods (Supplementary Data Figure 2). From the 19 clusters identified, 17 could be associated to specific cell lineages, while the remaining two were discarded. Based on the inclusion criteria described in the methods, 179304 out of 242224 cells (74.02%) were included for further analysis. Cell phenotypes were further clustered into functional groups using a set of functional markers. The functional clustering resulted in a total number of 47 functional cell populations (Figure 3A).

**Figure 3.**
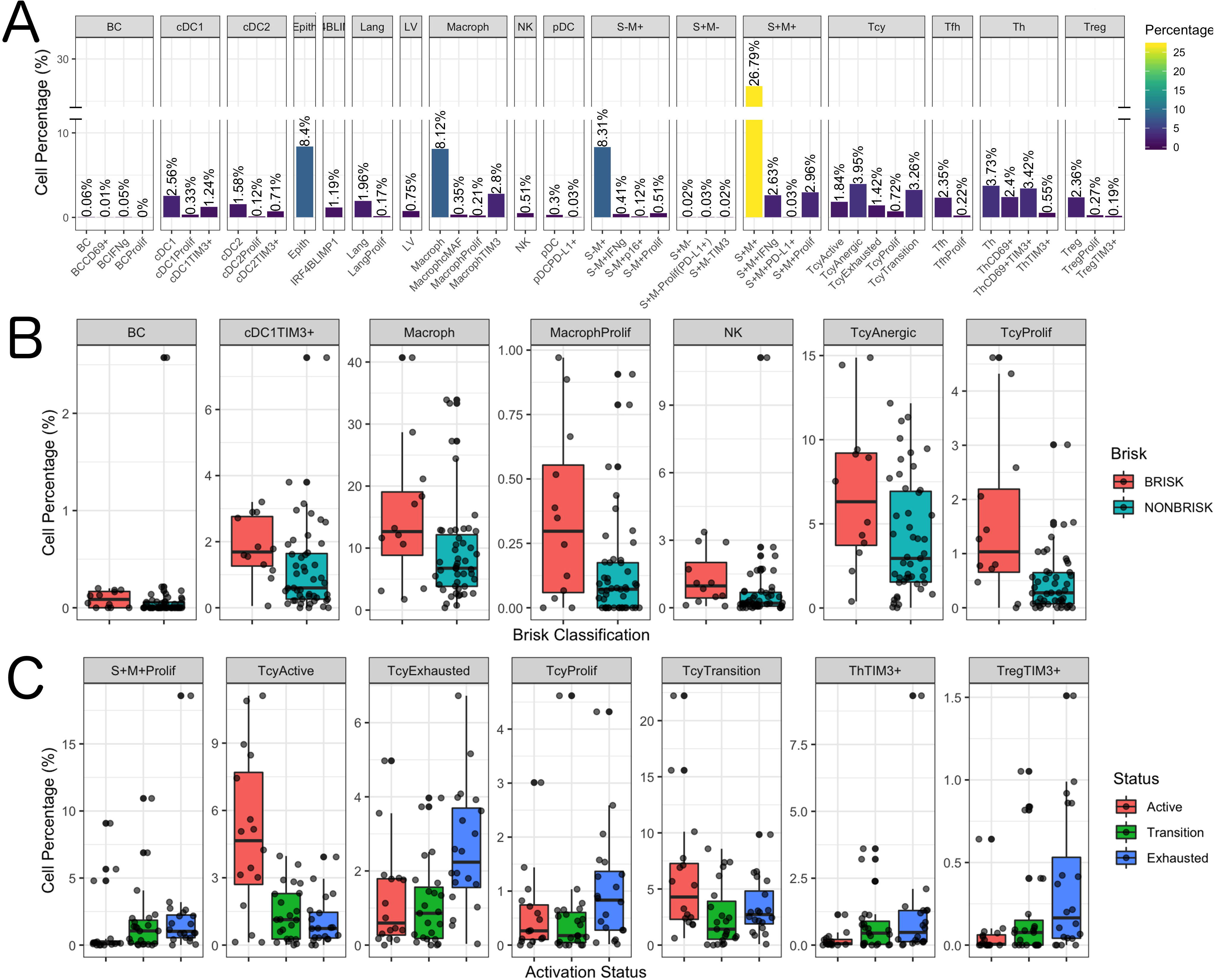
Distribution of the immune cells’ subgroups and significant differences in the morphological and functional TILs categories. The percentage of cells for each inflammatory subpopulation across all the cores is showed in (A). The 17 phenotypic clusters are on the upper side of the graph, while at the bottom each of them is subdivided in the respective functional subclusters. BC = B cells; cDC1 = classical dendritic cells type 1; cDC2 = classical dendritic cells type 2; Epith = epithelial cells; IRF4BLIMP1 = plasma cells; Lang = Langerhans cells; LV = lymph vessels; Macroph = macrophages; pDC = plasmocytoid dendritic cells; S-M+ = S100+MelanA-melanoma cells; S+M− = S100-MelanA+ = melanoma cells; S+M+ = S100+MelanA+ melanoma cells; Tcy = cytotoxic T cells; Tfh = T follicular helpers; Th = T helpers; Treg = regulatory T cells; suffix: “prolif” = proliferating, IFNg = interferon gamma. B) Significant differences (p.value < 0.05) in cell percentages between brisk and non-brisk categories (Wilcoxon rank sum test). Significant differences (p.value < 0.05) in cell percentages across the functional groups: Active, Transition, Exhausted (Kruskal-Wallis rank sum test).

The most abundant cell population consisted of melanoma cells (41.76%). Most of the melanoma cells expressed both melanocytic markers Melan A and S-100, while two minor groups of melanoma cells had loss of expression of one of the two markers. The second most abundant cell type were the macrophages (“Macroph”, CD68+CD163+Lysozyme+HLA-DR+, 11.48%), one quarter of them expressing the immunosuppressive marker TIM3. Epithelial cells represented 8.4% of the population of our data set. The lymphoid compartment accounted for multiple subpopulations. Tcy (CD3+CD8+) and T helpers (“Th”, CD3+CD4+FOXP3-) were the most abundant subtypes, accounting respectively for 11.19% and 10.10% of all the cells, while regulatory T cells (“Treg”, CD3+CD4+FOXP3+) represented 2,82% of the cells. We interpreted the last T cell cluster as T follicular helpers (“Tfh”, CXCL13+PD1+, 2.57%) even though this cluster did not express the full Tfh phenotype. Within Tcy, we could identify the active (CD69+OX40+/−, 16,1% of all the lymphocytes), transition (balanced expression of both exhaustion and expression markers, 29,6%) and exhausted (high expression of LAG3 and/or TIM3, low/absent CD69 and OX40, 12,8%) functional subgroups. Moreover, we found a clonally expanding subgroup (6%), and one with low or absent expression of all functional markers, that we defined as “anergic” (34,9%) but that could represent also naïve T cells. Th could also be further divided according to their expression of activation and exhaustion markers (28,5% ThCD69+, considered active, 18,46% ThCD69+TIM3+, considered transitional, and 13,08% ThTIM3+, considered immunosuppressive). NK cells, as expected in melanoma, were extremely infrequent (0.51%), but even more infrequent were the B cells (“BC”, CD20+), that represented 0.12% of the total. The CD20 negative cells characterized by high expression of IRF4 and Blimp1 (“IRF4_Blimp1”, 1,19%) were interpreted as plasma cells^13^. Finally, we could identify among the dendritic cell group the classical dendritic cells type 1 (“cDC1”, CD141+CD4+IRF8+, 4.13%), classical dendritic cells type 2 (“cDC2”, CD1c+CD4+HLA-DR+, 2.41%), Langerhans cells (“Lang”, CD1a+Langerin+, 2.13%), and plasmacytoid dendritic cells (“pDC”, CD123+, 0.33%). In both the two classical dendritic cell subgroups an immunosuppressive TIM3+ subpopulation was identifiable, while in the pDC group a small subpopulation was found to express PD-L1. No immunosuppressive subpopulation was identified among the Langerhans cells. Some subpopulations were statistically significantly different (p-val < 0.05) comparing brisk vs non-brisk and active vs transition vs exhausted. Brisk cases were significantly enriched in BC (p-val = 0.041), TIM3+cDC1 (p-val = 0.024), macrophages (p-val = 0.043) (including the proliferating subgroup (p-val = 0.034)), NK (p-val = 0.011), anergic Tcy (p-val = 0.039) and proliferating Tcy (p-val = 0.006) (Figure 3B). Active cores had, together with more active Tcy (p-val < 0.001), higher percentages of Tcy in transition (p-val = 0.030), transition cores more Th (p-val = 0.057), while exhausted cases had more TIM3+Tregs (p-val = 0.018) and more proliferating melanoma cells (p-val = 0.045) (Figure 3C).

### Neighbourhood analysis

We applied neighbourhood analysis in order to systematically identify social networks of cells and draw conclusions on actual cell-cell interactions. Macrophages and epithelial cells were in general most often located in strict proximity to the melanoma cells, without differences among the functional or morphologic categories. Brisk cases showed more Tcys in close proximity to melanoma cells than non-brisk cases, as expected (Supplementary Data Figure 3). Interestingly, brisk cases had a higher prevalence specifically of transition and active Tcy in contact with melanoma cells compared with non-brisk cases (Active/Exhausted ratio: Brisk = 2.108762, Non-Brisk = 1.331195), that instead had relatively more exhausted and anergic cells in contact with melanoma cells (Active/Anergic ratio: Brisk = 1.743081, NonBrisk = 0.7704947).

To understand what is the immune context that determines activation and exhaustion, we compared the results of neighbourhood analysis between active (Figure 4 A) and exhausted (Figure 4 B). The detection of the interaction of Langerhans cells with the epithelium, identifiable in both functional groups, was considered as positive control and proof of concept for the method. Some other interactions were present in both functional groups: TIM3+ macrophages and exhausted/translational Tcy, CD69+TIM3+Th and translational Tcy, active Th and anergic Tcy, TIM3+cDC1 and CD69+TIM3+Th. Cell-cell interactions that were specific for active cases included: active Th and active Tcy, TIM3+macrophages and CD69+TIM3+Th, NK and Tfh, Tfh and exhausted Tcy, and pDC and Langerhans. On the other hand, cell-cell interactions specific for exhausted cases included: CD69+TIM3+Th and anergic Tcy, anergic Tcy and active Th, anergic Tcy and BC expressing INFgamma, CD69+TIM3+Th and TIM3+cDC2, TIM3+macrophages and transition Tcy, NK and PD-L1+pDC, and the different subtypes of Tfh and cDC1.

**Figure 4.**
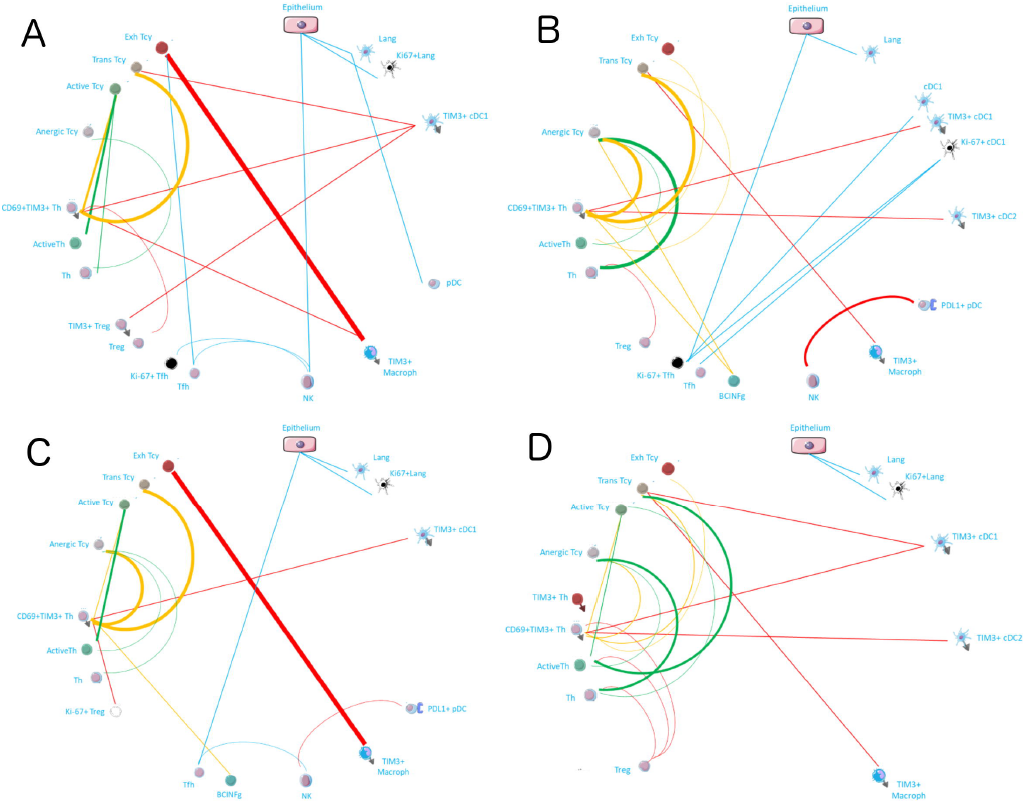
Neighbourhood analysis in the morphological and functional TILs categories. The result of the neighbourhood analysis for active (A), exhausted (B), brisk (C), and non-brisk (D) cases represented as cellular social networks. The thickness of the edge in the network represents the level of interaction between the different cell types. The colour of the line indicates interactions leading to immune suppression (red), to immune stimulation (green), to a probably sub-optimal/impaired immune stimulation (orange), no immune implications (blue).

### qPCR and shotgun proteomics

To confirm the existence of the same functional subcategories in metastatic melanoma samples, that represents the type of lesions that receive the immunotherapeutic treatment we perform qPCR followed by proteomics on microdissected TILs, comparing brisk and non-brisk cases using an “absent” case and a “tumoral melanosis” (i.e. complete melanoma regression with persistence of melanin-loaded macrophages) case as controls. Furthermore, we could measure directly the levels of expression of IFNg, the best indicator of CD8+ activation, for which no suitable antibody exists for FFPE material. 4/8 melanomas with a brisk TILs pattern, and 3/7 with non-brisk TILs pattern proved active, confirming that we could subclassify from a functional point of view also metastatic lesions (Supplementary Data Figure 4).

To correlate the gene expression measured by qPCR with the actual protein expression we performed a proteomic analysis on the microdissected material in 3 representative cases, i.e. one with high IFNg expression (“IFNg-high”), one with the high LAG3 and no IFNg expression (“LAG3-high”), and one with none of the four markers strongly expressed (“none”). The results showed a higher number of proteins identified in the “IFNg-high” sample (324) compared to “LAG3-high” (93) and “none” (134), with almost two thirds of them (210/324) not shared with the other two samples and enriched in proteins involved in different inflammatory pathways (innate immunity, TNFR1 signalling pathway, FAS signalling pathway, T cell receptor and Fc-epsilon receptor signalling pathway), including the interferon gamma-mediated signalling pathway. As we minimized the difference in the number of microdissected cells in each sample, these observations could only be explained by a higher production of pro-inflammatory proteins in the “IFNg-high” sample.

## Discussion

The fact that not all brisk infiltrates have a good prognosis^5,14^, that TILs populations can be very heterogeneous^15,16^ and that anti-PD-1 immunotherapy was found to support functionally activated T cells^6^ urged a thoughtful investigation of the functional status of the TILs. Since the molecules that mediate exhaustion are expressed upon activation in order to prevent the hyperactivity of the immune system^17^, to investigate the activation status of lymphocytes the simultaneous expression of several molecules must be evaluated at a single cell level and at once, as a panel^18^. Moreover, the assessment of the interactions between cells requires preservation of the tissue architecture. Our study is the first to assess up to 40 markers on tissue sections at single cell resolution without losing their spatial distribution. We obtained by high-dimensional in-situ immunotyping a snapshot of the co-expression patterns of activation and inhibition markers in tissue sections. OX40 was strongly correlated with Ki-67 and PD-1, confirming its role in sustaining Tcy clonal expansion^19^. LAG3 is expressed very early and only transiently during T cell activation, but persistent T cell activation with sustained expression of LAG3 together with other exhaustion markers (e.g. PD-L1) results in T cell dysfunction^20^. LAG3+Tregs have been shown to suppress DC maturation^21^.

We could observe that both brisk and non-brisk cases can harbour predominantly exhausted TILs or predominantly active TILs; therefore, with the morphological classification (brisk – non brisk – absent), a complementary functional classification (active – transitional - exhausted) co-exists. The importance of going beyond the morphologic classification of TILs was previously raised by others, who claimed that a functional analysis could help in the application of a better immunoscore for therapeutic prediction^16^. Only a minority (15%) of patients presented with all active cores, and 30% of patients presented with at least one active core, whereas the great majority of the patients presented with only exhausted or transition areas at the moment in which the melanoma was removed. Since the percentage of patients that obtains a durable response with single agent checkpoint therapy^22,23,24^ lies between 15 and 30%, and since Krieg et al^6^ reported that anti-PD-1 immunotherapy supports functionally activated T cells, it is tempting to speculate that the “mostly active” TILs cases could correspond to the responders to immunotherapy, while the “mostly exhausted” cases could benefit instead from combination approaches with immunotherapy and other type of therapies in order to rescue the exhausted T cells. Therefore, adding the functional evaluation could definitely improve the predictive value of the morphological TILs patterns in melanoma. Since spontaneous melanoma regression, present in only 10-35% of the melanomas, is considered to be the end-results of the melanoma-eliminating capacities of active TILs, we also studied the association of activation of TILs with early and late regression^25^. Our data showed a clear-cut association of activation of TILs with late regression areas, indirectly proving the functional meaning of an active infiltrate.

We obtained a functional picture of the inflammatory landscape in brisk versus non-brisk cases and in active/transition/exhausted TILs microenvironments. A previous study already casted some doubts on the significance of the brisk pattern, as they found no evidence of clonally expanded TILs in some cases with a brisk infiltrate^16^, and hypothesized that clonally expanded T cells might represent not only cytotoxic cells but also regulatory cells. We indeed found not only immune stimulating cells such as proliferating and anergic Tcy, cDC1, and NK significantly increased in brisk cases, but also immune suppressive cells such as BC and macrophages. Active and exhausted cases were instead functionally coherent categories. An active microenvironment was enriched for active Tcy, while an exhausted microenvironment was enriched not only with Treg, possible origin of the exhaustion, but also with proliferating melanoma cells, possible effect of the lack of immune control over the neoplasia.

A slight increase in Treg between active and exhausted cases may not be able alone to justify the shift from an active to an exhausted microenvironment. To this end, adding the spatial information and making inferences about the interactions between the cells in the tissue using neighbourhood analysis could represent a better way to investigate this dynamic. In this way, we could identify the most important differences that could explain the transition from an active to an exhausted environment. In active cases, it was possible to identify a stronger spatial association between active Th and active Tcy, while the same association was weaker in transition cases and disappeared in exhausted ones. In general, there was a decrease of interactions between the Th compartment and the active Tcy from active to exhausted cases. This could be explained observing the peculiar interactions with the other cell types in each of the functional states. In active cases, TIM3+ macrophages interacted with CD69+TIM3+ Th, as well as cDC1 expressing TIM3 interacted with TIM3+ Treg, possibly limiting their suppressive effect on the Tcy. In transition cases instead TIM3+ cDC1 are interacting only with CD69+TIM3+ Th and transitional Tcy, their block on Treg is removed and the Treg population is globally inhibiting all the Th subpopulation, probably reducing the strength of activation of the Tcy compartment. In fact, in active and exhausted cases the active Th are seen more often in contact with transitional Tcy, that are also the direct target of Tregs, while CD69+TIM3+ Th are in direct connection with exhausted Tcy. Starting from the transitional status we see the appearance of interactions between BC expressing INFg-related molecules and CD69+TIM3+ Th and anergic Tcy, and between TIM3+ cDC2 and CD69+TIM3+ Th, confirming the immune suppressive role of this cell types. The presence of a shift of interaction of the active Th towards anergic Tcy observed in the exhausted cases may be a rescue mechanism, meant to induce new active Tcy starting from anergic/naïve Tcy but possibly contrasted by the effects of the surrounding immunosuppressive cells. Finally, in active cases there is a strong interaction between TIM3+ macrophages and exhausted/transitional Tcy along all the statuses, confirming their prominent role in keeping the exhaustion in the tumoral microenvironment.

Considering the brisk and non-brisk classification, instead, we can see that both categories are a mixture of immune stimulating and suppressive interactions, once again confirming that the morphological categories are functionally very heterogeneous (Figure 4B). Paradoxically, it looks like non-brisk cases have more activating interactions than brisk cases. Nevertheless, the real determinant that may justify the good prognosis of the brisk cases may be that in brisk case melanoma cells are more in contact specifically with active/transition Tcy than non-brisk cases, that instead had more exhausted and anergic cells in contact with melanoma cells.

In conclusion, in this paper we have shown that the activation status of the TILs does not necessarily parallel the morphological categories, and that within a single melanoma, the inflammatory response may vary considerably. The classification of primary melanomas based on the functional paradigm had an improved prognostic value than that of the brisk classification. We hypothesize that the general good prognosis of melanomas with a brisk pattern of TILs could be based on the fact that the Tcy that are in contact with the melanoma cells at the moment that the melanoma is excised are still active, and consequently the melanoma is still under immune control. We have shown a bioinformatic pipeline that, starting from common immunofluorescence stainings, can transform the tissue into a digitized image, which represents the starting point for multiple deeper levels of analysis (Figure 5). In this study we have also described the main interactions between the inflammatory subpopulations in an active, transition and exhausted environment, interactions that should be taken into consideration when assessing the response to immunotherapy and that will ultimately lead to the identification of functional inflammatory microenvironments that may benefit from personalized combined therapy protocols.

**Figure 5.**
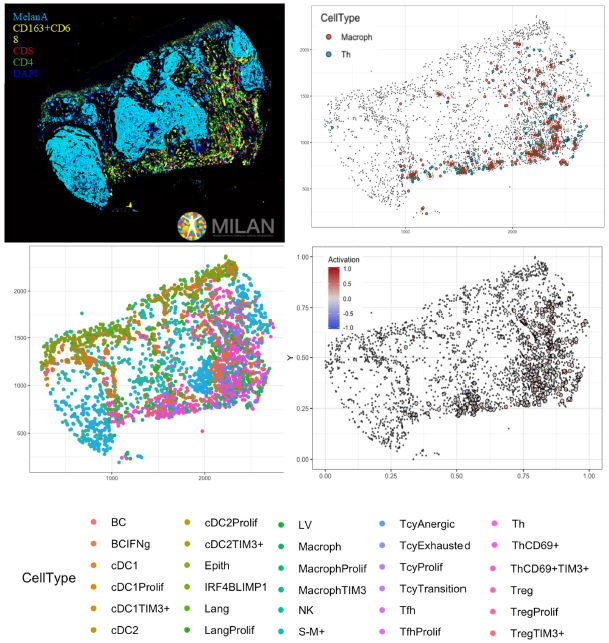
Complete digitalization of a core and practical example of a possible downstream analysis. Anti-clockwise: (1) images for the markers stained with the MILAN multiplexing technique (for this work 39 markers, but we reached in our laboratory an output of around 100 markers per single section) are acquired and composite images with some selected markers can be prepared. (2) All the markers are used to phenotypically identify all the cell subtypes present in the tissue and the tissue is digitally reconstructed. (3) Studies of the functional status of these cells can be done, for example we could localize exhausted and active cells in the tissue. (4) With neighbourhood analysis it is possible to identify the cells that are in proximity with each other more than chance can suggest, inferring possible interactions between these cell types (in the image, two cells identified as “neighbours” are encircled in red).

## Material and methods

### Sample Description

Twenty-nine invasive primary cutaneous melanomas from the Department of Pathology of the University Hospitals Leuven (KU Leuven), Belgium, were classified based on the H&E staining according to the pattern of the inflammatory infiltrate into brisk (6 cases) and non-brisk (23 cases). According to their subtype, 24 superficial spreading melanomas, 3 nodular melanomas, and 2 lentigo maligna melanoma were included.

### TMA construction

Tissue Micro Arrays (TMAs) were constructed with the GALILEO CK4500 (Isenet Srl, Milan, Italy). For each patient, one to five representative regions of interest were sampled according to the size of the specimen and the morphological heterogeneity both of the melanoma and of the distribution of the infiltrate. According to the morphological TILs pattern, we sampled brisk areas (i.e. 6 brisk areas in melanomas with brisk TILs pattern, termed “brisk in brisk” and 10 brisk areas in melanomas with non-brisk TILs pattern, termed “brisk in non-brisk”) and non-brisk areas (i.e. 15 non-brisk areas in non-brisk melanomas, termed “non-brisk in non-brisk”); in addition, areas showing “early regression” (7),” late regression” (5) and “no regression” (17) according to current morphological criteria^15^ were sampled. After processing, cutting and staining of the TMA blocks, a total of 60 cores were available for analysis.

### Multiplex-Stripping Immunofluorescence

The multiplex-stripping method was performed according to the MILAN technique, as published in Cattoretti G, et al^11^. The antibodies used for the multiplex staining are listed in the Table 2. Slides were scanned, and images acquired with the Hamamatsu Nanozoomer S60 scanner (Nikon, Italia) equipped with a Nikon 20X/0.75 lambda objective. Complete removal of previous layers was monitored by a) checking for unexpected staining consistent with the subcellular and tissue distribution of a previously detected marker; and b) spurious co-clustering of unrelated molecules in the hierarchical clustering images.

**Table 2.** Multiplex antibody panel description.

### Image Pre-processing

Fiji/ImageJ (version 1.51u) were used to pre-process the images (File format: from. ndpi to.tif 8/16 bit, grayscale). Registration was done through the Turboreg and MultiStackReg plugins, by aligning the DAPI channels of different rounds of staining, saving the coordinates of the registration as Landmarks and applying the landmarks of the transformation to the other channels. Registration was followed by autofluorescence subtraction (Image process → subtract), previously acquired in a dedicated channel, for FITC, TRITc and Pacific Orange. A macro was written in Fiji/ImageJ and used for the TMA segmentation into single images. Cell segmentation, mask creation, and single cell measurements were done with a custom pipeline using CellProfiler (version 2014-07-23T17:45:00 6c2d896). Quality Control (QC) over the Mean Fluorescence Intensity (MFI) values was performed using feature and sample selection. In short, those cells that did not have expression in at least three markers, and those markers that were not expressed in at least 1% of the samples were removed. MFIs were further normalized to Z-scores as recommended in Caicedo JC, et al^13^. Z-scores were trimmed between −5 and +5 to avoid a strong influence of any possible outliers in the downstream analysis. The correlation between the different markers was calculated using Pearson’s correlation coefficient.

### Functional analysis of TILs

We selected two activation (CD69, OX40) and two exhaustion (TIM3, LAG3) markers after literature review and preliminary testing of the antibody performance on control FFPE under the conditions of the multiplex protocol. The expression levels of these markers were measured selectively on CD8+ lymphocytes using a first mask focused only on CD8+ cells. Principal Component Analysis (PCA) was applied over the expression values to evaluate the functional structure of the data and to assign an activation value in the [−1, 1] range to each cell (Supplementary Information 1, Supplementary Data Figure 5). Briefly, principal components (PCs) 2 and 3 were used as the rotation matrix revealed that PC1 contained all the markers in the same direction (same sign). The point of maximum activation (Activation = 1), was defined where the projected value of CD69 (marker of activation) over PCs 2 and 3 was at the maximum while the point of maximum exhaustion (Activation = −1) where the projected value of TIM3 (marker of exhaustion) over PCs 2 and 3 was at the maximum (Figure 2). The gradient of transition was defined between the previously defined points and the centroid of the projected dataset. Pairwise t-tests with pooled standard deviation (sd) were used to find significant differences in the level of activation of the images regarding multiple histopathologic parameters (brisk/non-brisk infiltrate, regression, number of lymphocytes, ulceration, Breslow thickness, mitoses, subtype of melanoma). For this purpose, the activation level of each image was represented by the mean of its cells while the intrinsic degree of heterogeneity was captured by the sd. Continuous values of the histopathological parameters were fitted using linear models instead. P-values were adjusted for multiple comparisons using the holm method. A cut-off of 0.05 was used as significance threshold for the adjusted p-values. Images were further classified into: ‘Active’, ‘Transition’, and ‘Exhausted’ (from now on, indicating the core status) using one-tailed t-tests comparing the distribution of the activation values in a specific image versus the background distribution (combination of all images). P-values were adjusted using the False Discovery Rate (fdr) method. A cut-off value of 0.001 over the adjusted p-values was used as classification threshold.

For the survival analysis, the functional status of each patient was represented by the average level of activation of its cells and dichotomized into active (average level of activation > 0) and exhausted (average level of activation <= 0).

### Phenotypic identification

To evaluate the cell subpopulations, a second mask based on the DAPI nuclear staining contour expanded by 5 pixels was created. A two-tier approach was followed for the identification of cell subpopulations: phenotypic and functional. The phenotypic identification was conducted by applying three different clustering methods: PhenoGraph, ClusterX, and K-means, over the phenotypic markers: CD3, CD20, CD4, HLA-DR, Bcl6, CD16, CD68, CD56, CD141, CD1a, CD1c, Blimp1, Langerin, Lysozyme, Podoplanin, FOXP3, S100AB, IRF4, IRF8, CD1a, CXCL13, CD8, CD138, CD123, PD-1, and MelanA. PhenoGraph and ClusterX were implemented using the cytofkit package from R^12^. Clusters were represented by a vector containing the mean of each marker and were used to further associate them to a cell subpopulation using prior knowledge. For a phenotype to be assigned to a cell, at least two clustering methods should agree on their predicted phenotype. Prior to functional identification, PCA was repeated over the Tcy cells (CD8+) using CD69, OX40, LAG3, and TIM3 markers in order to confirm that with the new mask, the same dataset with the same structure as with the CD8+ mask could be retrieved (Supplementary Data Figure 1). The functional identification was conducted by applying PhenoGraph over the functional markers: CD69, Ki-67, TAP2, GBP1, MYC, p16, MX1, OX40, c-Maf, PD-L1, LAG3, TIM3, and Phospho-Stat1 (with the exception of Tcy cells for which we used a personalized panel consisting of: CD8, CD69, OX40, LAG3, TIM3, PD-1, and Ki-67). Clusters were represented, associated to cell subpopulations, and evaluated for stability as described for the phenotypic identification. Significant differences in the cellular composition of the cores based on activation status were identified using Kruskal-Wallis rank sum test. The same approach was repeated for the brisk infiltrate histopathologic parameter using Mann-Whitney test.

### Neighbourhood analysis

An unbiased quantitative analysis of cell-cell interactions was performed using an adaptation of the algorithm described in Schapiro D, et al^26^ for neighbourhood analysis to systematically identify social networks of cells and to better understand the tissue microenvironment. Our adaptation also uses a kernel-based approach (radius = 30 px) to define the neighbourhood of a cell and a permutation test (N = 1000) to compare the number of neighbouring cells of each phenotype in a given image to the randomized case. This allows the assignment of a significance value to a cell-cell interaction representative of the spatial organization of the cells. Significance values were further classified into avoidance (−1), non-significant (0), and proximity (1) using a significance threshold of 0.001 (more significant that all the random cases). Interactions across images were integrated according to equation 1:

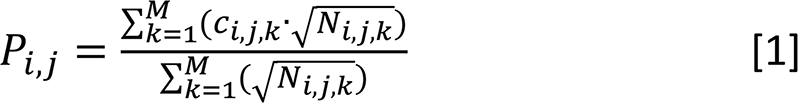

where *Ci,j,k* is the significance value (−1, 0, or 1) of the interaction between cell types *i* and *j* for image *k*, and *Ni,j,k* is the geometric average of the number of cells of type *i* and *j* for image *k*. Cell-cell interactions were considered strong if they were significant in at least 75% of the N-adjusted cases (abs(P) > 0.75), moderate if 50%, (0.5 < abs(P) <= 0.75), weak if 25% (0.25 < abs(P) <= 0.5), and non-significant otherwise (abs(P) <= 0.25). A comparative analysis of the above described method was performed for the different core statuses as well as for the different brisk infiltrate cases (Figure 4).

Even though neighbourhood analysis allows evaluation of cell-cell interactions, the mathematical model applied is limited in cases where there are dominant cell types that grow in nests, with cells packed next to each other. Therefore, for melanoma cells, we evaluated the closest neighbours by counting the number of cells of each subpopulation (apart from melanoma cells) that were in their neighbourhood and divided the amount by the geometric average of the number of melanoma cells and the number of cells of the specific population across all the cores in which the specific population appears. This analysis was repeated for the different brisk and activation cases.

### Laser Microdissection

18 fresh frozen melanoma metastases with different types of TILs patterns (9 brisk, 7 non brisk, 1 absent and 1 tumoral melanosis) were collected in the Department of Pathology of the University Hospitals Leuven (KU Leuven), Belgium. 10 micrometre-thick sections were cut from each fresh frozen block and put on a film slide (Zeiss, Oberkochen, Germany). Sections were stained with crystal violet. Areas with dense TILs infiltrate were microdissected with the Leica DM6000 B laser microdissection device (Leica, Wetzlar, Germany). A calculation was made in order to microdissect the same surface in all the samples in order to minimize differences between the samples (around 10.000 lymphocytes/sample).

### qPCR of laser microdissected samples

RNA extraction was done with RNeasy^®^ Plus Micro Kit (Qiagen) according to the protocol. cDNA retrotranscription followed by an amplification step was done with Ovation^®^ Pico SL WTA System V2 (Nugen) according to protocol. Primers for Interferon gamma (INFg, forward ‘TGTTACTGCCAGGACCCA’ and reverse ‘TTCTGTCACTCTCCTCTTTCCA’), TIM3 (forward ‘CTACTACTTACAAGGTCCTCAGAA’ and reverse ‘TCCTGAGCACCACGTTG’), LAG3 (forward ‘CACCTCCTGCTGTTTCTCA’ and reverse ‘TTGGTCGCCACTGTCTTC’), CD40-L (forward ‘GAAGGTTGGACAAGATAGAAGATG’ and reverse ‘GGATAAGGATCTTTCTCCTGTGTT’), CD45 (forward ‘GCTACTGGAAACCTGAAGTGA’ and reverse ‘CACAGATTTCCTGGTCTCCAT’), Beta2microglobulin (forward ‘ACAGCCCAAGATAGTTAAGTG’ and reverse ‘ATCTTCAAACCTCCATGATGC’), HPRT (forward ‘ATAAGCCAGACTTTGTTGGA’ and reverse ‘CTCAACTTGAACTCTCATCTTAGG’) were designed with Perl Primer^®^ and tested in our laboratory. 96-wells plates were loaded with Fast SYBR^®^ Green Master Mix, the primers and the samples in the recommended proportions, and analysed with the 7900 HT Fast Real-Time PCR system (Applied Biosystems). The log fold change (logFC) of the expression values towards the expression value of CD45 were calculated. If the log(IFNg/CD45) was positive, the sample was classified as positive. On the other hand, exhaustion was defined by expression of LAG3 and/or TIM3 with lack of IFNg and CD40L expression.

### Shotgun Proteomics of laser microdissected samples

The materials used for the shotgun proteomics analysis were: Trifluoroacetic acid, MS grade porcine trypsin, DTT (dithiothreitol), IAA (Iodoacetamide), ABC (Ammonium Bicarbonate), HPLC grade water, acetonitrile (ACN), were from Sigma-Aldrich (Sigma-Aldrich Chemie GmbH, Buchs, Switzerland). All solutions for Mass Spectrometry (MS) analysis were prepared using HPLC-grade. LCM collected material corresponding to about 10^4 cells for each sample group was re-suspended in 90 μl of bidistilled water and immediately stored at −80°C. For the bottom-up MS analysis, all the samples were processed and trypsinized. Briefly, thawed cells were submitted to a second lysis adding 60 μl of 0.25% w/v RapiGest surfactant (RG, Waters Corporation) in 125mM ammonium bicarbonate (ABC) and sonicated for 10 min. Samples were then centrifuged at 14 000 × g for 10 min. About 140 μl of supernatants were collected, transferred in a new tube and quantified using bicinchoninic acid assay (Pierce -Thermo Fisher Scientific). After 5min denaturation (95°C), proteins were reduced with 50mM DTT in 50mM ABC at room temperature and alkylated with 100mM IAA in 50mM ABC (30 min incubation in dark). Digestion of samples was performed overnight at 37°C using 2μg of MS grade trypsin. RG surfactant were removed using an acid precipitation with TFA at a final concentration of 0.5% v/v. Samples were then spun down for 10 min at 14000 × g and supernatants collected for MS analysis. Peptide mixtures were desalted and concentrated using Ziptip™ μ-C8 pipette tips (Millipore Corp, Bedford, MA). An equal volume of eluted digests was injected at least three times for each sample into Ultimate™ 3000 RSLCnano (ThermoScientific, Sunnyvale, CA) coupled online with Impact HD™ UHR-QqToF (Bruker Daltonics, Germany). In details, samples were concentrated onto a pre-column (Dionex, Acclaim PepMap 100 C18 cartridge, 300 μm) and then separated at 40°C with a flow rate of 300 nL/min through a 50 cm nano-column (Dionex, ID 0.075mm, Acclaim PepMap100, C18). A multi-step gradient of 4 hours ranging from 4 to 66% of 0.1% formic acid in 80% ACN in 200min was applied^27^. NanoBoosterCaptiveSpray™ ESI source (Bruker Daltonics) was directly connected to column out. Mass spectrometer was operated in data-dependent acquisition mode, using CID fragmentation assisted by N2 as collision gas setting acquisition parameters as already reported^28^. Mass accuracy was assessed using a specific lock mass (1221.9906 m/z) and a calibration segment (10 mM sodium formate cluster solution) for each single run. Raw data from nLC ESI-MS/MS were elaborated through DataAnalysis™ v.4.1 Sp4 (Bruker Daltonics, Germany) and converted into peaklists. Resulting files were interrogated for protein identification through in-house Mascot search engine (version: 2.4.1), as described^28^. Identity was accepted for proteins recognized by at least one unique and significant (p-value < 0.05) peptide.

### Pathways Analysis

Gene-set enrichment analysis was performed with DAVID 6.8^29,30^. Pathways were visualized and partially analysed with STRING v10^31^.

## Acknowledgements

Francesca Maria Bosisio is funded by the MEL-PLEX research training programme (‘Exploiting MELanoma disease comPLEXity to address European research training needs in translational cancer systems biology and cancer systems medicine’, Grant agreement no: 642295, MSCA-ITN-2014-ETN, Project Horizon 2020, in the framework of the MARIE SKŁODOWSKA-CURIE ACTIONS).

Asier Antoranz is funded by the SyMBioSys research training programme (‘Systematic Modeling of Biological Systems’), grant agreement no: 675585, MSCA-ITN-2015-ETN, Project Horizon 2020, in the framework of the MARIE SKŁODOWSKA-CURIE ACTIONS.

Maddalena Maria Bolognesi is supported by a clinical research project BEL114054 (HGS1006-C1121) of the University of Milano-Bicocca and GlaxoSmithKline, UK.

---

**Supplementary Figure 1 - Biplot showing the projection of the Tcy cells over PCs 2 and 3.** In order to verify the solidity of the method, we compared the results obtained with the DAPI mask with the ones obtained with the CD8+ mask (Figure 2.A). We applied the same approach to define the activation status of the T cytotoxic cells on the Tcy phenotypic cluster, and we observed the same structural behaviour of the activation and exhaustion markers (CD69 and TIM3 at opposite ends of the activation spectrum, concordance between LAG3 and OX40).

**Supplementary Figure 2. Phenotype Identification.** A two-tier approach was implemented for the identification of cell subpopulations. A) Initially, KMeans, PhenoGraph, and ClusterX (left) were used to identify the main inflammatory subpopulations, resulting in clusters that were named by manual annotation by the pathologist based on the level of expression of the phenotypic markers (right). The final phenotypes were defined using a consensus-based approach (a phenotype was assigned if two or more clustering methods agreed in their prediction). B) Cell populations were further sub-clustered into functional subgroups using PhenoGraph over the set of functional markers and annotation by the pathologist.

**Supplementary Figure 3 - Analysis of the interactions between melanoma cells and the inflammatory subpopulations.** With this alternative approach of neighbourhood analysis, we observed that the main inflammatory cells subtypes in contact with melanoma cells are macrophages and (as expected) epithelial cells, both in brisk and non-brisk cases, followed by Tcy with active and in transition Tcy in brisk cases and proliferating and anergic Tcy in non-brisk cases. Other small differences between brisk and non-brisk cases are more TIM3+ cDC1, cDC2 and TIM3+ macrophages in contact with melanoma cells in brisk cases.

**Supplementary Figure 4 - qPCR and shotgun proteomics.** Confirmation of the functional subgroups by qPCR (A) and shotgun proteomics plus pathway analysis (B). With qPCR we were able to identify, both in B and NB metastasis, active (green rectangles) and exhausted (red rectangles) cases. With proteomics we confirmed that the “IFNg-high” case had more proteins that the “LAG3-high” and the “none” case involved in inflammatory pathways among which the IFNg-related pathway (in grey).

**Supplementary Figure 5 - Definition of activation.** The rotation matrix from the PCA shows that PC1 is capturing general expression of the initial markers (all the markers with the same sign). On the other hand, in PC2, TIM3 (exhaustion) has a different sign than the rest of the markers, and the same is true for PC3 and CD69 (activation). Therefore, PC2 and PC3 were used for the definition of the activation function.

## Supplementary Data

The followed pipeline is detailed below: PC2 and PC3 are mapped into polar coordinates.

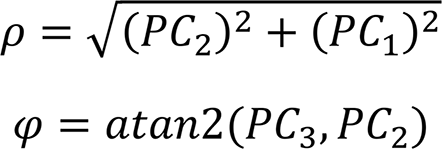

where PC2 and PC3 are calculated from the rotation matrix

PC2 = 0.0444 · CD69 + 0.7048 · OX40 + 0.4764 · LAG3 – 0.5236 · TIM3

PC3 = −0.7505 · CD69 + 0.3656 · OX40 + 0.1196 · LAG3 + 0.5372 · TIM3

The point of maximum activation (Activation = 1) was defined as the point where the projected value of CD69 in PCs 2 and 3 reaches a maximum (Supplementary Data Figure 6, point A). The angle corresponding to the multi-valued inverse tangent of the rotation vectors of PC3 and PC2 (atan2(PC3, PC2)) (*φ*0) is added to *φ*.

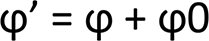

The point of maximum exhaustion (Activation = −1) was defined as the point where the projected value of TIM3 in PCs 2 and 3 reaches a maximum (Supplementary Data Figure 6, point B).

The line of transition (Activation = 0) was defined as the bisector between the projected vectors of LAG3 and OX40 over PCs 2 and 3 (Supplementary Data Figure 6, Transition Line).

The 4 resulting areas (supplementary figure 1, 1 to 4) do not cover the same range of *φ*. Each area was scaled so that it covers 90 degrees (π/2 rads).

Finally, the value of activation of each cell was calculated as:

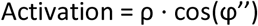

where ρ is the radius and *φ*” the scaled angle.

## Notes

#### Summary of Updates

The pipeline has been improved. The results from neighbourhood analysis were further detailed. Prognostic information has been added.

