## Supplementary figures and images for "Functional heterogeneity of lymphocytic patterns in primary melanoma dissected through single-cell multiplexing"

### Supplementary data figure 1

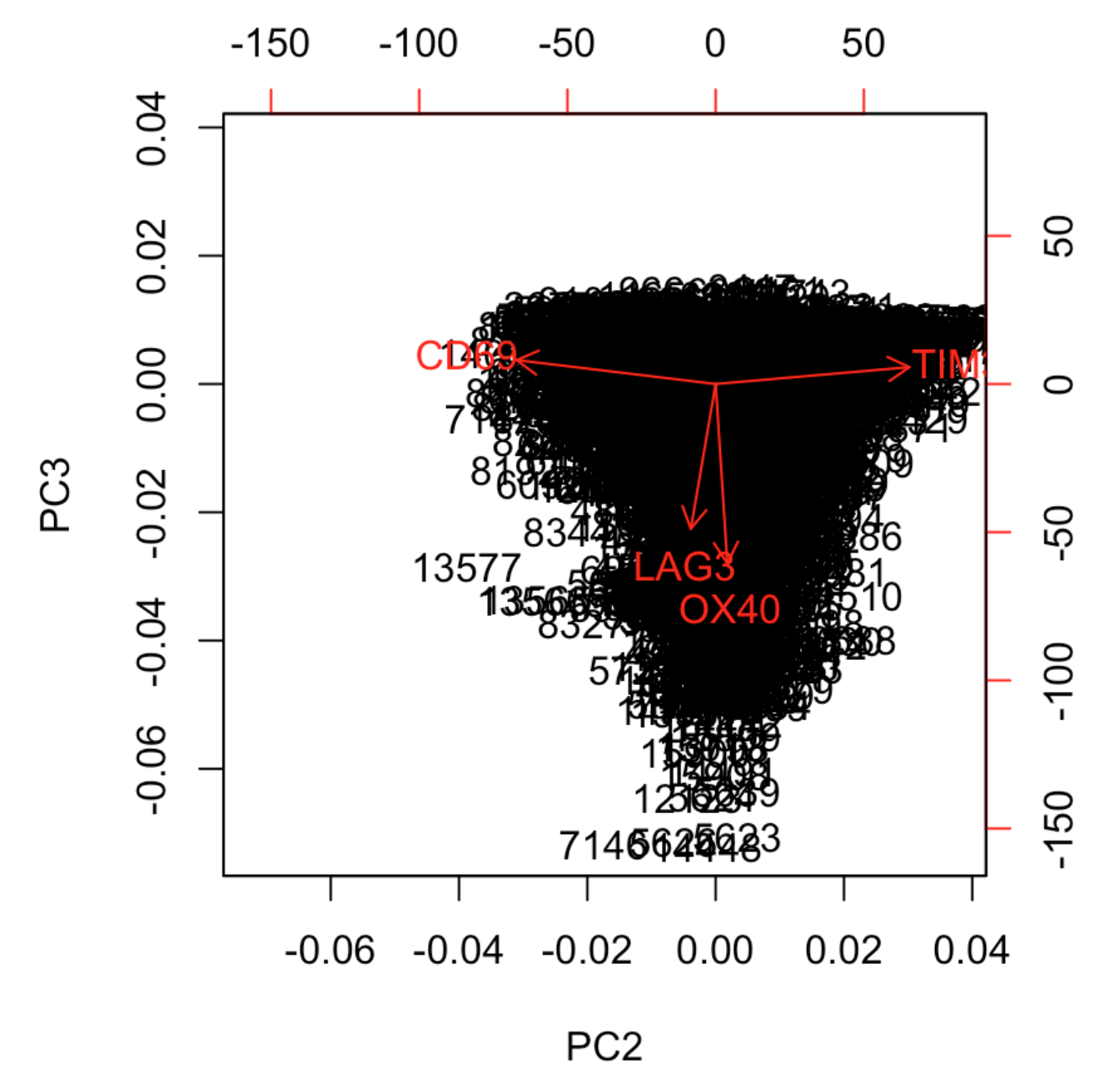

### Supplementary data figure 2

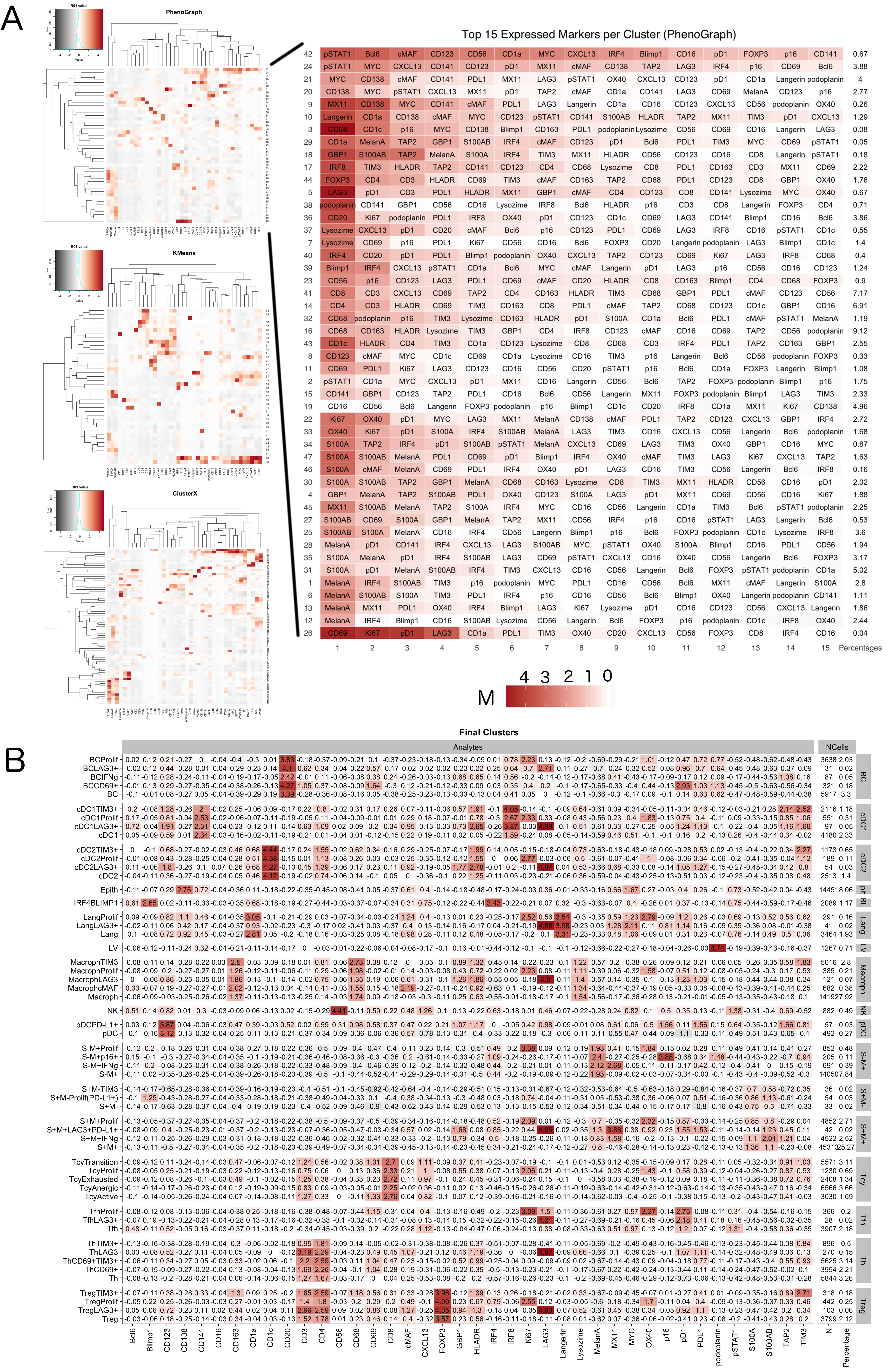

### Supplementary data figure 3

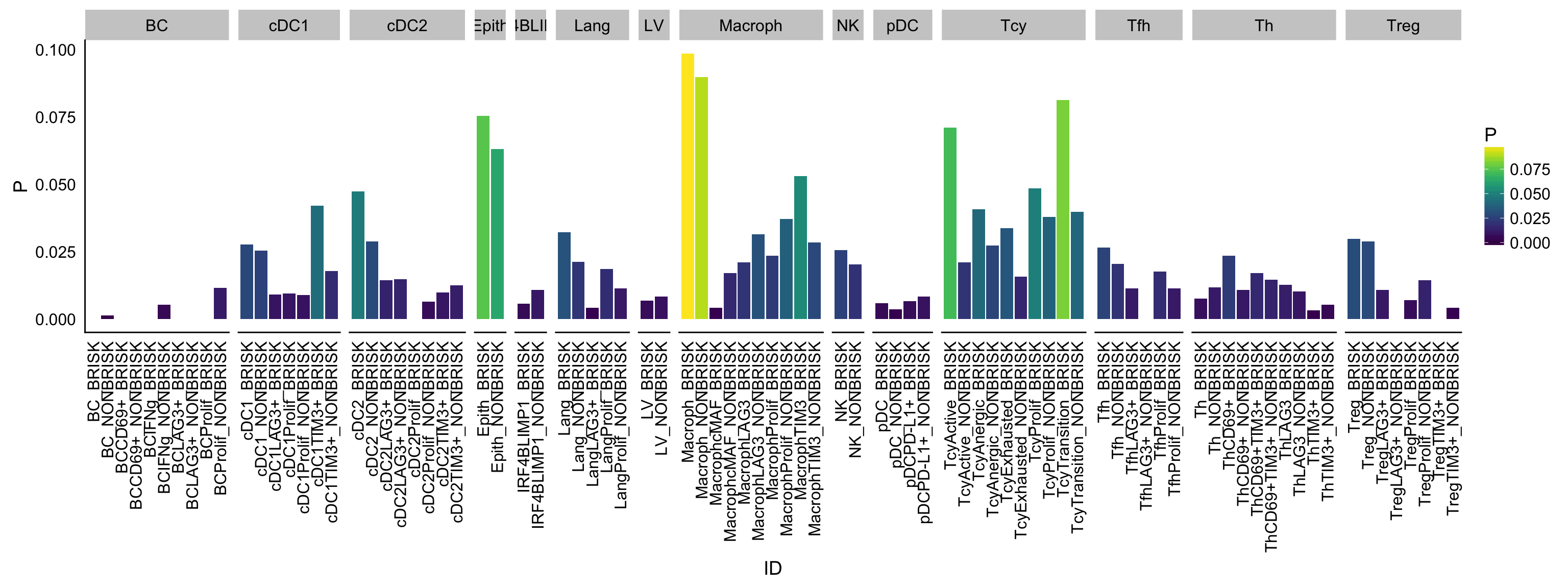

### Supplementary data figure 4

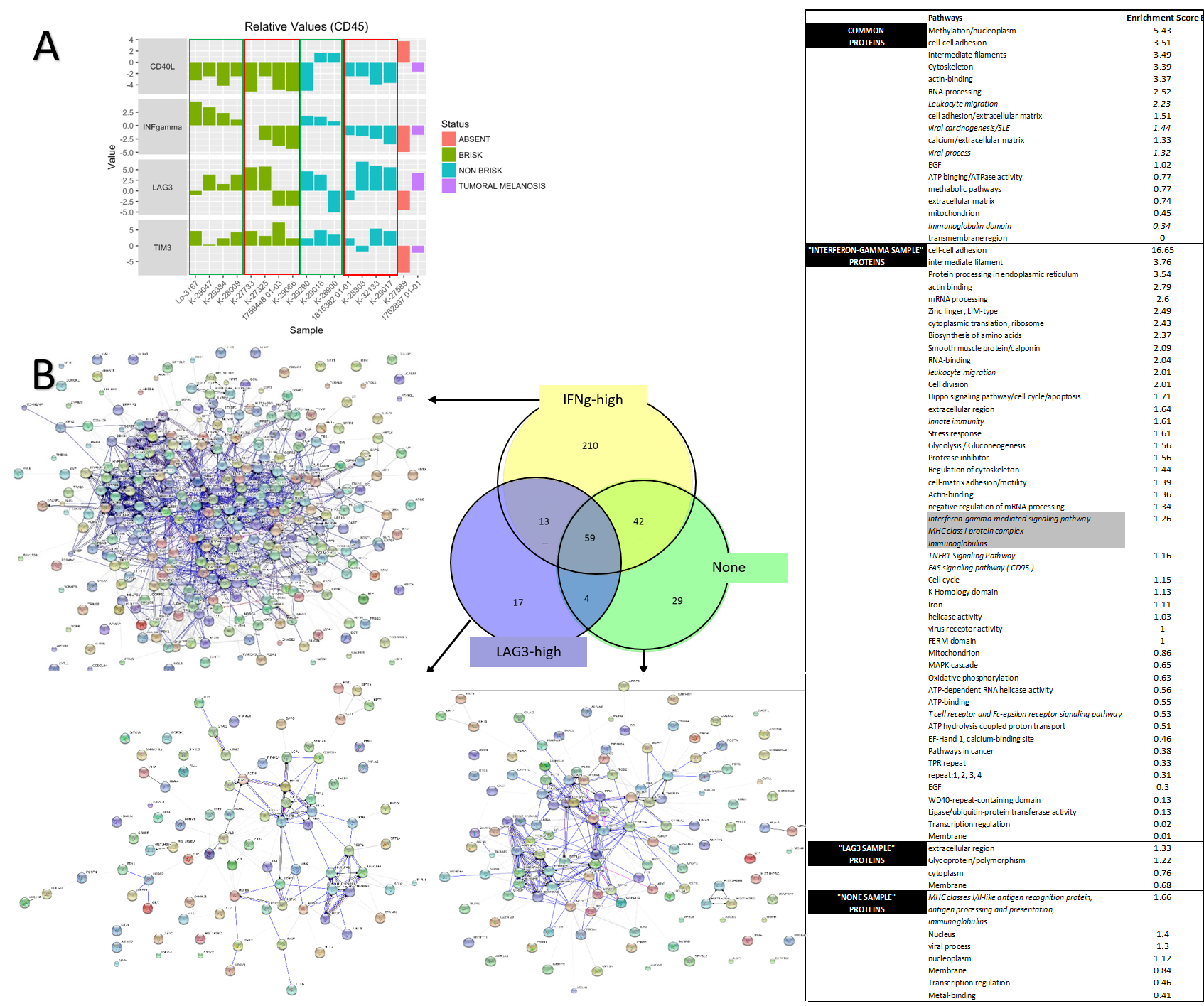

### Supplementary data figure 5

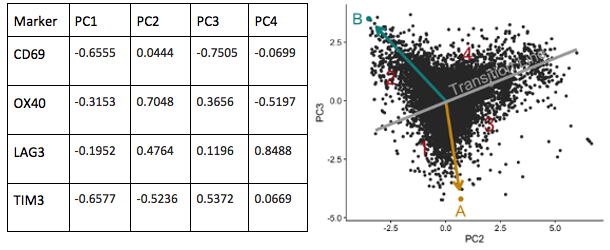
